# Dynamics of brain function in chronic pain patients assessed by microstate analysis of resting-state electroencephalography

**DOI:** 10.1101/2020.09.29.318246

**Authors:** Elisabeth S. May, Cristina Gil Ávila, Son Ta Dinh, Henrik Heitmann, Vanessa D. Hohn, Moritz M. Nickel, Laura Tiemann, Thomas R. Tölle, Markus Ploner

**Affiliations:** Department of Neurology, School of Medicine, Technical University of Munich (TUM), Munich, Germany; TUM-Neuroimaging Center, School of Medicine, TUM, Munich, Germany; Center for Interdisciplinary Pain Medicine, School of Medicine, TUM, Munich, Germany

**Author notes:** Correspondence to: Markus Ploner, Department of Neurology, Technical University of Munich (TUM), Ismaninger Str. 22, 81675 Munich, Germany. Both authors contributed to this work equally (Elisabeth S. May, Cristina Gil Ávila).

**Keywords:** chronic pain, dynamics, EEG, microstate analysis, resting-state

## Abstract

Chronic pain is a highly prevalent and severely disabling disease, which is associated with substantial changes of brain function. Such changes have mostly been observed when analyzing static measures of brain activity during the resting-state. However, brain activity varies over time and it is increasingly recognized that the temporal dynamics of brain activity provide behaviorally relevant information in different neuropsychiatric disorders. Here, we therefore investigated whether the temporal dynamics of brain function are altered in chronic pain. To this end, we applied microstate analysis to eyes-open and eyes-closed resting-state electroencephalography (EEG) data of 101 patients suffering from chronic pain and 88 age- and gender-matched healthy controls. Microstate analysis describes EEG activity as a sequence of a limited number of topographies termed microstates, which remain stable for tens of milliseconds. Our results revealed that sequences of 5 microstates, labelled with the letters A to E, described resting-state brain activity in both groups and conditions. Bayesian analysis of the temporal characteristics of microstates revealed that microstate D has a less predominant role in patients than in healthy participants. This difference was consistently found in eyes-open and eyes-closed EEG recordings. No evidence for differences in other microstates was found. As microstate D has been previously related to attentional networks and functions, abnormalities of microstate D might relate to dysfunctional attentional processes in chronic pain. These findings add to the understanding of the pathophysiology of chronic pain and might eventually contribute to the development of an EEG-based biomarker of chronic pain.

## 1. Introduction

Chronic pain is a highly disabling disease, which affects 20-30% of the adult population [7; 24]. Its pathophysiology is not fully understood and treatment is often insufficient [59], imposing a tremendous burden on patients, health care systems and society [46]. Converging lines of evidence have shown that chronic pain is associated with extensive changes of brain structure and function [2; 29]. Understanding these changes promises fundamental insights into the underlying pathophysiology and might eventually help to establish a much sought-after biomarker of chronic pain [14; 57].

Brain function in chronic pain has mostly been assessed using functional magnetic resonance imaging (fMRI) [2] and electro-/magnetoencephalography (EEG/MEG) [44]. Most of these studies have analyzed static measures of brain activity during the resting-state, usually by aggregating a certain feature of brain function across several minutes. However, brain activity varies over time and it is increasingly recognized that these temporal dynamics provide behaviorally and clinically relevant information, which complements static measures [19; 45]. Correspondingly, it has been proposed that the dynamics of brain activity and connectivity critically shape the perception of pain [28]. By assessing brain activity and connectivity at ultra-low frequencies below 0.1 Hz, recent fMRI studies have provided support for this concept in chronic pain (e.g. [3; 5; 10; 58]). However, the temporal dynamics of chronic pain-related brain activity at frequencies higher than 1 Hz have not been consistently explored yet.

EEG and MEG are well suited to study such dynamic changes of brain activity at higher frequencies. One of the best-established methods in this field is microstate analysis (see [25; 34] for reviews), which has revealed that temporal changes of EEG activity do not occur randomly. Instead, EEG activity switches between a limited number of so-called microstates. During a microstate, the EEG topography remains stable for tens of milliseconds before abruptly transitioning to another microstate. EEG resting-state activity is usually well-described with 4 to 6 microstates, which are remarkably similar across participants. Thus, microstate analysis quantifies resting-state EEG recordings as sequences of a limited number of microstates. The temporal characteristics of these microstates carry important information about mental processes [6; 34]. Moreover, abnormalities of temporal microstate characteristics have been observed in different neuropsychiatric disorders (e.g. [12; 38; 48]). During the writing of this manuscript, a first microstate study in patients suffering from chronic pain was published. The results showed lower occurrence and time coverage of microstate C in patients with fibromyalgia [20]. However, these findings need to be replicated and extended to other chronic pain conditions.

Here, we investigated whether the temporal dynamics of brain activity are changed in a large cohort of patients suffering from chronic pain. To this end, we applied microstate analysis to EEG resting-state recordings of 101 patients suffering from different types of chronic pain and 88 matched healthy control participants. Thereby, the study aimed to further the understanding of the pathophysiology of chronic pain and to potentially contribute to the development of a brain-based biomarker of chronic pain.

## 2. Materials and methods

### 2.1. Participants

The current study represents a re-analysis of previously published data obtained at the Technical University of Munich for the large-scale study of brain dysfunction in chronic pain [54]. 101 patients (69 women; age = 58.1 ± 13.6 years [mean ± SD]) suffering from different types of chronic pain and 88 age- and sex-matched healthy controls (60 women, age = 57.5 ± 14.2 years) participated in the study. Inclusion criteria for patients were a clinical diagnosis of chronic pain, with pain lasting at least 6 months, and a minimum reported average pain intensity of at least 4 out of 10 during the past 4 weeks (0 = no pain, 10 = worst imaginable pain). Exclusion criteria for patients were acute changes of the pain condition during the past 3 months (e.g. due to recent injuries or surgeries), major neurological diseases (e.g. epilepsy, stroke or dementia), major psychiatric diseases aside from depression and severe general diseases. Patients taking benzodiazepines were also excluded. Other medication was not restricted and was maintained. In total, 47 patients with chronic back pain, 30 patients with chronic widespread pain, 6 patients with joint pain and 18 patients with neuropathic pain were included in the study. Exclusion criteria for healthy participants were a past medical history of pain lasting more than 6 months, having any pain on the day of testing, surgery or acute injury during the last 3 months and any neurological or psychiatric diseases. All participants provided written informed consent. The study was approved by the ethics committee of the Medical Faculty of the Technical University of Munich and conducted according to the relevant guidelines and regulations.

Questionnaires were used to assess pain characteristics and comorbidities on the day of the recording. All patients completed the following questionnaires: pain characteristics were assessed by the short-form McGill pain Questionnaire (SF-MPQ) [33], depression by the Beck Depression Inventory II (BDI-II) [4] and anxiety by the State-Trait Anxiety Inventory (STAI) [52]. The medication was quantified for all patients using the Medication Quantification Scale (MQS) [21]. 81 patients additionally completed the painDETECT questionnaire [18] to assess the neuropathic pain component and 47 patients completed the Pain Disability Index (PDI) [16] and the Veteran’s RAND 12-Item Health Survey (VR-12) [50] to assess pain disability and quality of life, respectively. All healthy control participants completed BDI-II and STAI questionnaires to assess potential comorbidities. Detailed characteristics of the participants can be found in Table 1.

**Table 1.** Demographic data and questionnaire results. BDI = Beck Depression Inventory; STAI = State-Trait Anxiety Inventory; SF-MPQ = Short-form McGill Pain Questionnaire; VAS = Visual Analogue Scale; Avg. pain intensity = average pain intensity in the past 4 weeks; PDI = Pain Disability Index; PDQ = painDETECT; VR-12 PCS = Veteran’s Rand 12 Physical Component Summary; VR-12 MCS = Veteran’s Rand 12 Mental Component Summary; MQS = Medication Quantification Scale. Please note that data for avg. pain intensity in the past 4 weeks, pain duration and PDQ were only available for a subset of 81 patients. Data from PDI, VR-12 PCS and VR-12 MCS were only available for a subset of 47 patients.

| | Chronic pain patients<br>(mean ± SD) | Healthy controls<br>(mean ± SD) |
| --- | --- | --- |
| Number | 101 | 88 |
| Gender (m/f) | 32/69 | 28/60 |
| Age (years) | 58.2 ± 13.5 | 57.5 ± 14.3 |
| BDI | 15.8 ± 8.9 | 3.5 ± 4.5 |
| STAI - State | 39.5 ± 10.6 | 30.6 ± 6.1 |
| STAI - Trait | 44.0 ± 11.2 | 30.9 ± 7.1 |
| SF-MPQ total pain score | 27.1 ± 9.4 | – |
| Current pain intensity (VAS) | 5.2 ± 1.9 | – |
| Avg. pain intensity in the past 4 weeks (VAS) | 5.6 ± 1.6 | – |
| Pain duration (months) | 121.8 ± 114.4 | – |
| PDQ | 17.4 ± 6.5 | – |
| PDI | 27.4 ± 14.2 | – |
| VR-12 PCS | 31.8 ± 7.8 | – |
| VR-12 MCS | 46.4 ± 11.9 | – |
| MQS | 6.8 ± 8.1 | – |

### 2.2. Recordings

Brain activity was recorded using electroencephalography (EEG) during the resting-state. Participants were instructed to stay in a wakeful and relaxed state without performing any particular task. Two 5-min blocks of continuous resting-state data were recorded, one with eyes closed and the other with eyes open. The temporal order of the blocks was counter-balanced. During the recording, participants were comfortably seated and listened to white noise played through headphones to mask any ambient noise.

Data were recorded with 64 electrodes and a BrainAmp MR plus amplifier (Brain Products, Munich, Germany). The electrodes included all electrodes from the 10-20 international system and the additional electrodes Fpz, CPz, POz, Oz, Iz, AF3/4, F5/6, FC1/2/3/4/5/6, FT7/8/9/10, C1/2/5/6, CP1/2/3/4/5/6, TP7/8/9/10, P1/2/5/6/7/8 and PO3/4/7/8/9/10 (Easycap, Herrsching, Germany). Two electrodes were placed below the outer canthus of each eye to monitor eye movements. All EEG electrodes were referenced to FCz and grounded at AFz. For 81 patients and 69 healthy controls, muscle activity was simultaneously recorded with two bipolar electromyography (EMG) electrode montages and a BrainAmp ExG MR amplifier (Brain Products, Munich, Germany). EMG electrodes were placed on the right masseter and neck (semispinalis capitis and splenius capitis) muscles [13]. The EMG ground electrode was placed at vertebra C2. Data were obtained at a sampling frequency of 1000 Hz, with 0.1 μV resolution and were band-pass filtered online between 0.016 and 250 Hz. Impedances were kept below 20 kΩ.

### 2.3. Preprocessing

Preprocessing was performed with the Brain Vision Analyzer software (Brain Products, Munich, Germany) on the appended data from the eyes-open and the eyes-closed conditions. For artifact identification, a high pass filter at 1 Hz and a notch filter at 50 Hz were applied to remove low frequency drifts and electrical line noise, respectively. Independent Component Analysis was performed [23]. Components representing eye movements and muscle artifacts were identified based on their time courses and topographies and subtracted from the raw unfiltered EEG time series [60]. Signal jumps higher than ± 100 µV and their adjacent time intervals (200 ms before and after the jump) were marked for rejection. Subsequently, all data sets were visually inspected and remaining bad intervals were marked for rejection. Finally, data were re-referenced to the average reference and the reference electrode FCz was added to the electrode array.

### 2.4. Microstate analysis

Microstate analysis was performed using the free academic software Cartool version 3.8 (Brunet et al., 2011), Matlab (Mathworks, Natick, MA) and the Matlab toolbox Fieldtrip [41]. Analyses were performed separately for the eyes-open and eyes-closed conditions. For all analyses, the first minute of artifact-free continuous data of each condition was selected. Subsequently, the selected data segment was band-pass filtered between 1 and 40 Hz and down-sampled to 125 Hz, in line with previous studies (Britz, Van De Ville and Michel, 2010; Custo et al., 2017; Tomescu et al., 2018). Eyes-closed data of 1 chronic pain patient and 1 healthy control and eyes-open data of 11 patients and 8 controls had to be excluded from further analysis because no continuous one-minute artifact-free segment was available. Thus, final sample sizes were 100 patients and 87 healthy controls for the eyes-closed condition and 90 patients and 80 healthy controls for the eyes-open condition. An overview of the microstate analysis pipeline can be found in Figure 1.

**Figure 1.**
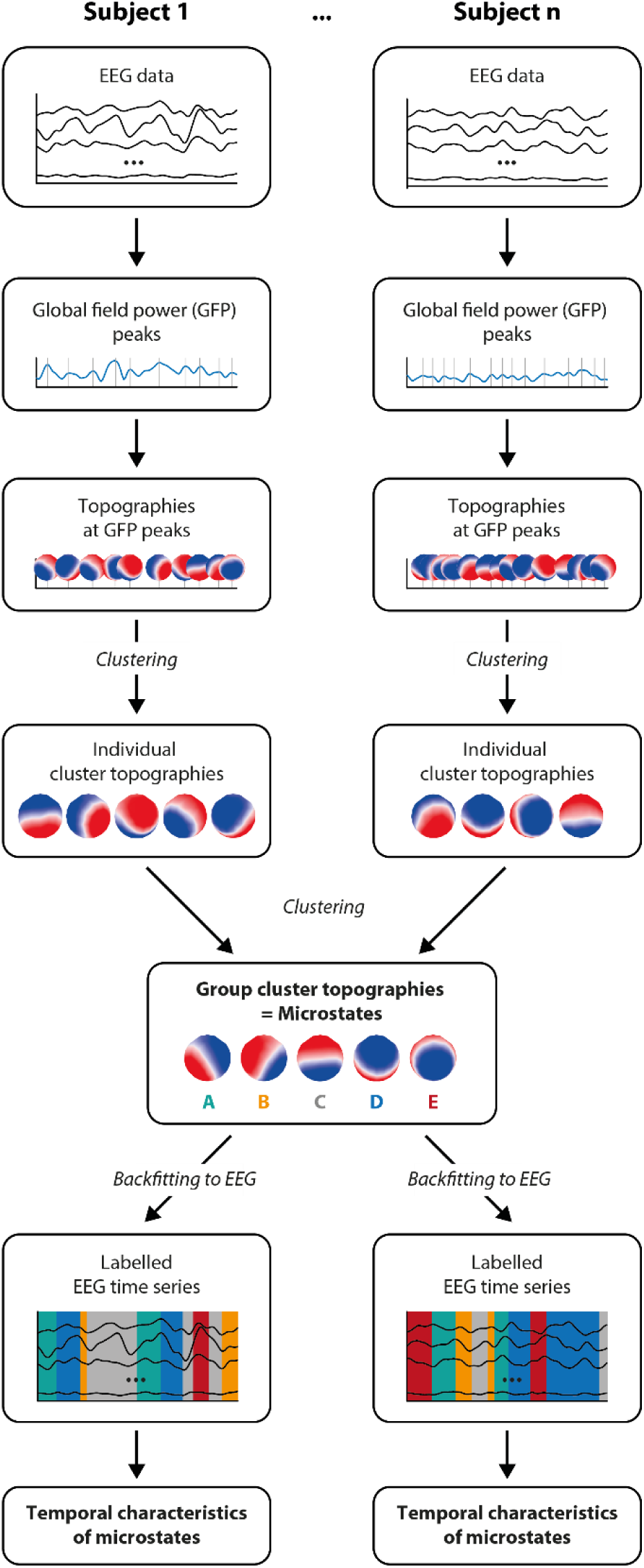
Microstate analysis. For each participant, the global field power (GFP) is calculated and topographies at GFP peaks are selected for individual clustering. Topographies at GFP peaks are clustered with a modified k-means clustering, leading to a variable number of individual cluster topographies per individual. Next, individual cluster topographies are concatenated and clustered on a group level. This resulted in 5 different group cluster topographies, labelled as microstates A to E. Microstate topographies are then fitted back to the individual EEG data, resulting in a labelled EEG time series in which each time point is associated to a microstate. From the labelled EEG time series, the temporal characteristics of microstates are derived. This analysis was performed separately per group (chronic pain patients and healthy participants) and condition (eyes-open and eyes-closed).

#### 2.4.1. Definition of microstates

We defined microstates through a well-established two-step clustering procedure using a modified k-means algorithm [42]. In line with previous studies [12; 38; 48; 55], this was done separately for each group and condition.

The first step consisted of a k-means clustering performed at the individual level. For each participant, EEG topographies at global field power (GFP) peaks were clustered, yielding a variable number of individual-level topographies. The GFP is a measure of the instantaneous strength of EEG activity measured over the whole scalp and mathematically defined as the standard deviation of the signals of all electrodes [39]. EEG topographies were clustered at GFP maxima since they represent the time points of highest signal-to-noise ratio [39; 42].

The clustering algorithm requires an a priori definition of k, which is the number of clusters into which the data will be grouped. To select the optimal number of clusters, we performed the clustering with different numbers of k = 4 to 12 initial clusters, following Cartool’s default settings for resting-state data. First, an initial number of k topographies was randomly selected from all GFP-peak topographies of the individual EEG time series. Second, the selected topographies were spatially correlated with the remaining topographies at GFP peaks, ignoring polarity. Third, the topographies at GFP peaks were assigned to the cluster with the highest spatial correlation. If the highest correlation was smaller than 0.5, the topography was not assigned to any cluster. Fourth, the center of each cluster was computed, resulting in k new “average” cluster topographies. The new cluster topographies were then again correlated with the topographies at GFP peaks, closing the loop. The algorithm stopped when the variance of the clusters converged to a limit. In order to overcome the random selection of the initial cluster topographies, the clustering was repeated 100 times per set of k clusters and the set explaining most variance of the data was selected. The optimal number of clusters was identified for each individual separately according to a meta-criterion with 7 independent optimization criteria (for more details refer to [6]). This procedure resulted in 4 to 8 topographies for each participant and condition. The number of individual topographies did not differ between groups for the eyes-open or the eyes-closed condition (eyes-closed: mean_patients_ = 5.020, mean_controls_ = 4.919, t = -0.727, p = 0.468, BF_10_ = 0.204, median δ = -0.099, 95% CI = [-0.377, 0.178]; eyes-open: mean_patients_ = 4.944, mean_controls_ = 4.912, t = -0.256, p = 0.799, BF_10_ = 0.171, median δ = -0.036, 95% CI = [-0.326, 0.253]; two-sided independent samples t-tests). In the second step, a second k-means clustering was performed at group level, clustering the concatenated individual topographies obtained in the previous step. Cartool’s default settings for the second clustering were adopted, i.e. an initial number of k = 4 to 15 clusters and 200 k-means initializations. Again, the polarity was ignored and a maximum correlation higher than 0.5 was needed for cluster assignment. The same meta-criterion as before was used to identify the optimal number of clusters on a group level. It consistently revealed an optimal number of 5 group-level topographies for each group and condition.

Each set of 5 group-level topographies (Figure 2) was visually inspected and compared to topographies reported in the literature. For both groups, the first four topographies closely resembled the four well-known ‘canonical’ microstates A to D reported previously and were labeled accordingly [25; 27; 34]. The topography of the fifth microstate closely resembled a microstate, which has been consistently reported in more recent studies [6; 11; 62], since an increasing number of studies is now using a data-driven approach to define the optimal number of microstates. We labeled it with the letter E. Throughout the manuscript, the 5 group-level topographies are referred to as microstates A to E. Similarities and differences of microstate topographies between groups were assessed by calculating spatial correlations and topographic analyses of variance (TANOVA) for all microstates (A to E), respectively. The spatial correlation is a scalar value computed as the Pearson’s correlation coefficient between all matched electrodes of two different topographies [39]. TANOVA is a non-parametric randomization test based on the global map dissimilarity of individual topographies [39]. The global map dissimilarity is a measure of the difference between two topographies directly related to the spatial correlation [39]. For each microstate, global map dissimilarity was computed between the microstate topographies of the patient and control groups using Cartool [9]. To obtain a p-value, this dissimilarity was compared to a distribution of dissimilarities, which was generated by randomly shuffling individual topographies between patient and control groups and re-computing the dissimilarity between the center topographies of the randomized groups. The process was repeated 5000 times. This comparison resulted in a p-value per microstate, which was given by the proportion of permutations in which the dissimilarity was smaller than the dissimilarity originally observed in the data. For both the eyes-closed and eyes-open conditions, resulting p-values were corrected for multiple comparisons across the 5 microstate classes using the False Discovery Rate (FDR) [61].

**Figure 2.**
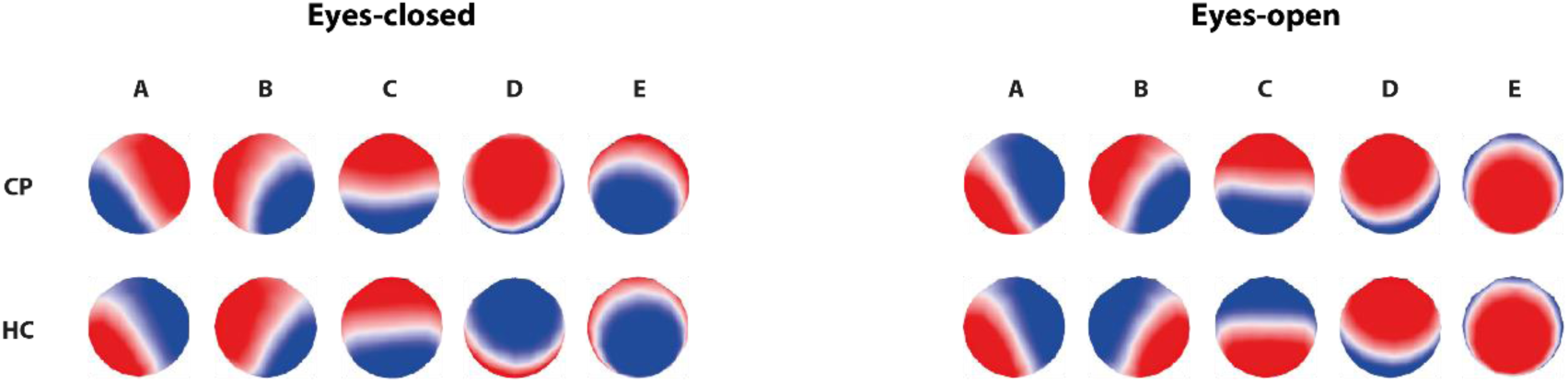
Microstate topographies of chronic pain patients and healthy participants for the eyes-closed and eyes-open conditions. Microstates were labelled with the letters A to E according to previous literature [34]. CP: Chronic pain; HC: Healthy control.

#### 2.4.2. Temporal microstate characteristics

Next, we determined the temporal characteristics of the five microstates for both groups. To this end, individual EEG time series were construed as time series of microstates through a ‘fitting procedure’, i.e. a microstate was assigned to every time point. For each participant, the EEG topographies of all time points were spatially correlated to the five microstate topographies of the participant’s group (patients/controls), ignoring polarity. Next, each EEG time point was assigned to a microstate (A to E). To ensure a certain continuity in the microstate time series, the relabeling was performed based on two criteria: (1) the correlation should be high and (2) the majority of the surrounding time points should belong to the same microstate [9]. To fulfill this compromise between goodness of fit and smoothness, standard temporal smoothing (window half size = 5 and strength (Besag factor) = 10) was applied [42; 55]. No label was assigned if the highest spatial correlation was smaller than 0.5. On average, the percentage of unlabeled time points was smaller than 0.2% and no differences existed between groups in any condition (eyes-closed: mean_patients_ = 0.077 %, mean_controls_ = 0.072 %, t = -0.308, p = 0.758, BF_10_ = 0.224, median δ = 0.166, 95% CI = [-0.319, 0.235]; eyes-open: mean_patients_ = 0.177 %, mean_controls_ = 0.106 %, t = -0.875, p = 0.383, BF_10_ = 0.2237, median δ = -0.124, 95% CI = [-0.416, 0.165]; two-sided independent samples t-tests).

Based on the time series of microstates, four measures were calculated to quantify the temporal characteristics of each microstate: mean duration, time coverage, frequency of occurrence and global explained variance (GEV). The mean duration is the average time (in milliseconds) for which a microstate persists before transitioning to a different microstate. The time coverage is the percentage of total time that a microstate is present. The frequency of occurrence is the number of times that a microstate recurs per second. The global explained variance is the percentage of global variance that is explained by every microstate.

Finally, we investigated the microstate sequence by examining the transition probabilities from each microstate to the others [31; 40; 55]. To this end, we computed the matrix of transition counts among all microstates for each participant and divided it by the overall count of transitions.

### 2.5. Statistical analysis

Group differences of temporal microstate measures (mean duration, time coverage, frequency of occurrence and global explained variance) and transition probabilities were analyzed in JASP version 0.13.1 [22] using two-sided independent sample t-tests in both frequentist and Bayesian frameworks. For the frequentist approach, significance level was set to 0.05. For p-values of temporal measures, FDR correction was performed in Matlab (Mathworks, Natick, MA) across the 5 microstates and the 4 different temporal measures, resulting in a correction for 20 statistical tests per condition (eyes-open/eyes-closed). For the transition matrix, FDR correction was performed across all 20 transitions per condition (eyes-open/eyes-closed). For the Bayesian analysis, default priors (Cauchy distributions with a scale parameter r = 0.707) were used. In addition to t-values and FDR-corrected p-values, results are reported using the two-tailed Bayes factor BF_10_. Effect size estimates for the BF_10_ are reported as the median of the posterior Cohen’s δ distribution together with its 95% credibility interval.

Finally, we investigated relationships between temporal microstate measures and clinical parameters, using JASP version 0.13.1 [22]. To this end, temporal microstate measures of microstate D (mean duration, time coverage, frequency of occurrence, global explained variance) were selected for a correlation analysis, since they consistently showed at least moderate evidence for a difference between patients and controls in both conditions in the previous analysis. Pearson’s correlations were calculated between the microstate measures and major clinical parameters, which were available for all patients (current pain intensity, SF-MPQ total pain score, depression (BDI) and medication (MQS)). Correlations were again calculated in both frequentist and Bayesian frameworks. In the Bayesian analysis, default priors (stretched beta priors with width = 1) were used. Results are reported using the Pearson’s correlation coefficient, its FDR-corrected p-value, its Bayes factor (BF_10_) and the 95% credibility interval of the correlation coefficient. FDR correction of p-values was performed in Matlab (Mathworks, Natick, MA) across all 16 performed correlations separately for each condition (eyes-open/eyes-closed).

### 2.6. Data and code availability

Raw and preprocessed EEG data in BIDS format [43] as well as scripts for statistical analyses are openly available at osf.io.

## 3. Results

The current study investigated whether the dynamics of resting-state brain activity are altered in patients suffering from chronic pain. We performed microstate analysis, which describes the time course of EEG activity as a sequence of a limited number of short stable topographies termed microstates. We applied microstate analysis to eyes-open and eyes-closed resting-state EEG activity and compared temporal characteristics of microstates between a large cohort of patients suffering from chronic pain and age- and sex-matched healthy control participants.

### 3.1. Definition of microstates A to E in patients and controls

We identified microstates using a standard two-step k-means clustering procedure [42]. The procedure consistently revealed 5 different microstates in both groups and conditions (Fig. 2). In accordance with previous studies [6; 8; 11; 26; 27; 34; 35; 62], the microstates were labeled as microstates A to E. Together, the five microstate explained 79.63% and 80.15% of the variance across individuals in the eyes-closed condition and 80.35% and 79.95% in the eyes-open condition (chronic pain patients and healthy controls, respectively), which is in good accordance with previous studies [12; 34; 49]. Figure 2 shows the topographies of both groups. The high similarity of topographies between groups was confirmed by high spatial correlations (eyes-closed: microstate A: r = 0.95, B: r = 0.93, C: r = 0.97, D: 0.89, E: r = 0.83; eyes-open: microstate A: r = 0.98, B: r = 0.99, C: r = 0.99, D: r = 0.96, E: r = 0.99). In addition, TANOVAs revealed subtle group differences between topographies hardly visible to the naked eye (eyes-closed: microstate A: p = 0.014, B: p < 0.001, C: p < 0.001, D: p < 0.001, E: p < 0.001; eyes-open: A: p = 0.005, B: p = 0.53, C: p < 0.001, D: p < 0.001, E: p = 0.31).

Taken together, the clustering procedures for both groups and conditions resulted in 5 microstate topographies, which were largely similar between groups.

### 3.2. Temporal characteristics of microstates in patients and controls

To investigate the dynamics of brain activity, we next analyzed whether the temporal characteristics of the 5 microstates differed between patients and healthy controls. To this end, microstates were back-fitted to the individual EEG time series by correlating microstate topographies with the EEG topographies at every time point. This allowed to assign each time point to a microstate, and, thus, to construe the EEG time series as time series of microstates.

We specifically calculated the mean duration, time coverage, frequency of occurrence and global explained variance of each microstate. This was done for each patient and each healthy control participant for the eyes-closed (Figure 3) and eyes-open (Figure 4) conditions. We next compared these temporal microstate characteristics between groups. The results revealed that all microstate D characteristics differed between patients and healthy participants. Such differences were consistently found in both conditions. In the eyes-closed condition, we found very strong evidence for a lower global explained variance of microstate D in patients compared to controls (Table 2; BF_10_ > 300, FDR-corrected p-value = 0.001). In addition, we found moderate evidence for a shorter mean duration, lower time coverage and lower frequency of occurrence of microstate D (Table 2; 3 < BF_10_ < 10, FDR-corrected p-values < 0.05). Similarly, in the eyes-open condition, the data provided moderate evidence for a shorter mean duration, lower time coverage, lower frequency of occurrence and lower global explained variance of microstate D in chronic pain patients (Table 3; 3 < BF_10_ < 10, FDR-corrected p-values < 0.05). For all other microstates, we did not find evidence for group differences consistent across measures and conditions (Tables 2 and 3).

**Table 2.**
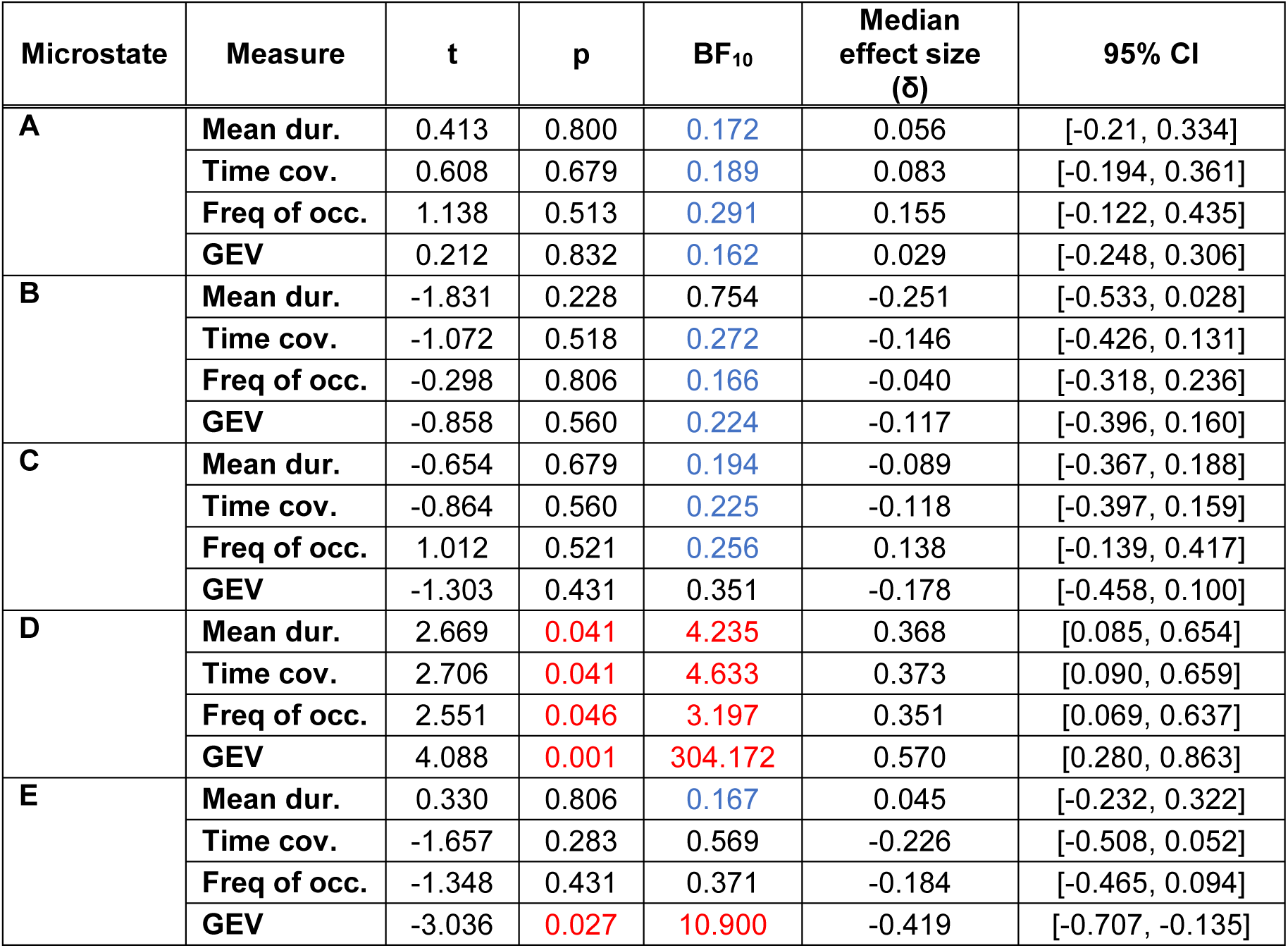
Group comparisons of temporal microstate measures in the eyes-closed condition. Results of two-sided independent samples t-test (frequentist and Bayesian approach). P-values are FDR-corrected. Color coded in red p < 0.05 and BF_10_ > 3, indicating at least moderate evidence for the alternative hypothesis and in blue BF_10_ < 1/3, indicating at least moderate evidence for the null hypothesis. Median effect sizes (δ) and their respective 95% credible interval (CI) are reported. Mean dur. = mean duration; Time cov. = time coverage; Freq of occ. = frequency of occurrence; GEV = global explained variance; BF_10_ = Bayes Factor in favor of the alternative hypothesis.

**Table 3.** Group comparisons of temporal microstate measures in the eyes-open condition. Results of two-sided independent samples t-test (frequentist and Bayesian approach). P-values are FDR-corrected. Color coded in red p < 0.05 and BF_10_ > 3, indicating at least moderate evidence for the alternative hypothesis and in blue BF_10_ < 1/3, indicating at least moderate evidence for the null hypothesis. Median effect sizes (δ) and their respective 95% credible interval (CI) are reported. Mean dur. = mean duration; Time cov. = time coverage; Freq of occ. = frequency of occurrence; GEV = global explained variance; BF_10_ = Bayes Factor in favor of the alternative hypothesis.

| Microstate | Measure | t | p | $BF_{10}$ | Median effect size ($\delta$) | 95% CI |
| --- | --- | --- | --- | --- | --- | --- |
| **A** | Mean dur. | -1.207 | 0.458 | 0.326 | -0.171 | [-0.465, 0.118] |
|  | Time cov. | 0.189 | 0.939 | 0.169 | 0.027 | [-0.262, 0.317] |
|  | Freq of occ. | 0.943 | 0.534 | 0.251 | 0.134 | [-0.156, 0.426] |
|  | GEV | 0.128 | 0.939 | 0.167 | 0.018 | [-0.271, 0.308] |
| **B** | Mean dur. | -1.051 | 0.603 | 0.224 | -0.114 | [-0.406, 0.175] |
|  | Time cov. | 0.098 | 0.939 | 0.167 | 0.014 | [-0.275, 0.304] |
|  | Freq of occ. | 1.317 | 0.421 | 0.370 | 0.187 | [-0.103, 0.481] |
|  | GEV | -0.726 | 0.624 | 0.212 | -0.103 | [-0.394, 0.186] |
| **C** | Mean dur. | -0.804 | 0.491 | 0.277 | -0.149 | [-0.442, 0.140] |
|  | Time cov. | -1.417 | 0.395 | 0.420 | -0.202 | [-0.496, 0.089] |
|  | Freq of occ. | -0.305 | 0.939 | 0.174 | -0.043 | [-0.333, 0.246] |
|  | GEV | -1.418 | 0.395 | 0.421 | -0.202 | [-0.496, 0.089] |
| **D** | Mean dur. | 2.828 | 0.026 | 6.402 | 0.407 | [0.110, 0.708] |
|  | Time cov. | 2.938 | 0.026 | 8.501 | 0.423 | [0.126, 0.725] |
|  | Freq of occ. | 2.942 | 0.026 | 8.595 | 0.424 | [0.127, 0.726] |
|  | GEV | 2.922 | 0.026 | 8.159 | 0.421 | [0.124, 0.723] |
| **E** | Mean dur. | -1.733 | 0.339 | 0.664 | -0.247 | [-0.543, 0.044] |
|  | Time cov. | -1.145 | 0.461 | 0.305 | -0.163 | [-0.455, 0.127] |
|  | Freq of occ. | -0.075 | 0.939 | 0.167 | -0.011 | [-0.300, 0.279] |
|  | GEV | -1.486 | 0.395 | 0.461 | -0.211 | [-0.506, 0.079] |

**Figure 3.**
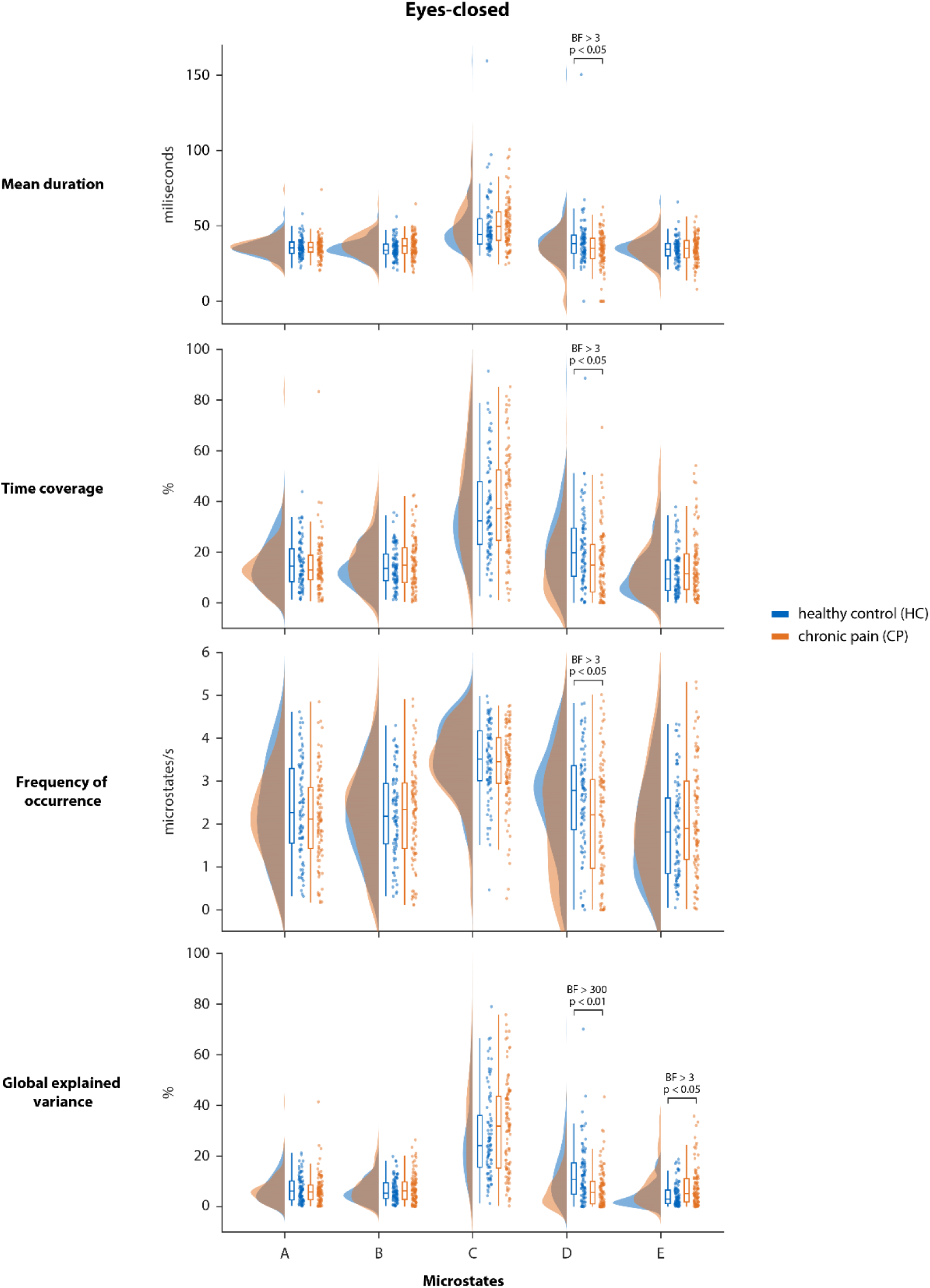
Temporal microstate characteristics in the eyes-closed condition. Mean duration, time coverage, frequency of occurrence and global explained variance of each microstate were calculated for both chronic pain patients and healthy controls. Raincloud plots [1] show un-mirrored violin plots displaying the probability density function of the data, boxplots, and individual data points. Boxplots depict the sample median as well as first (Q1) and third quartiles (Q3). Whiskers extend from Q1 to the smallest value within Q1 – 1.5* interquartile range (IQR) and from Q3 to the largest values within Q3 + 1.5* IQR. BF = Bayes Factor in favor of the alternative hypothesis.

**Figure 4.**
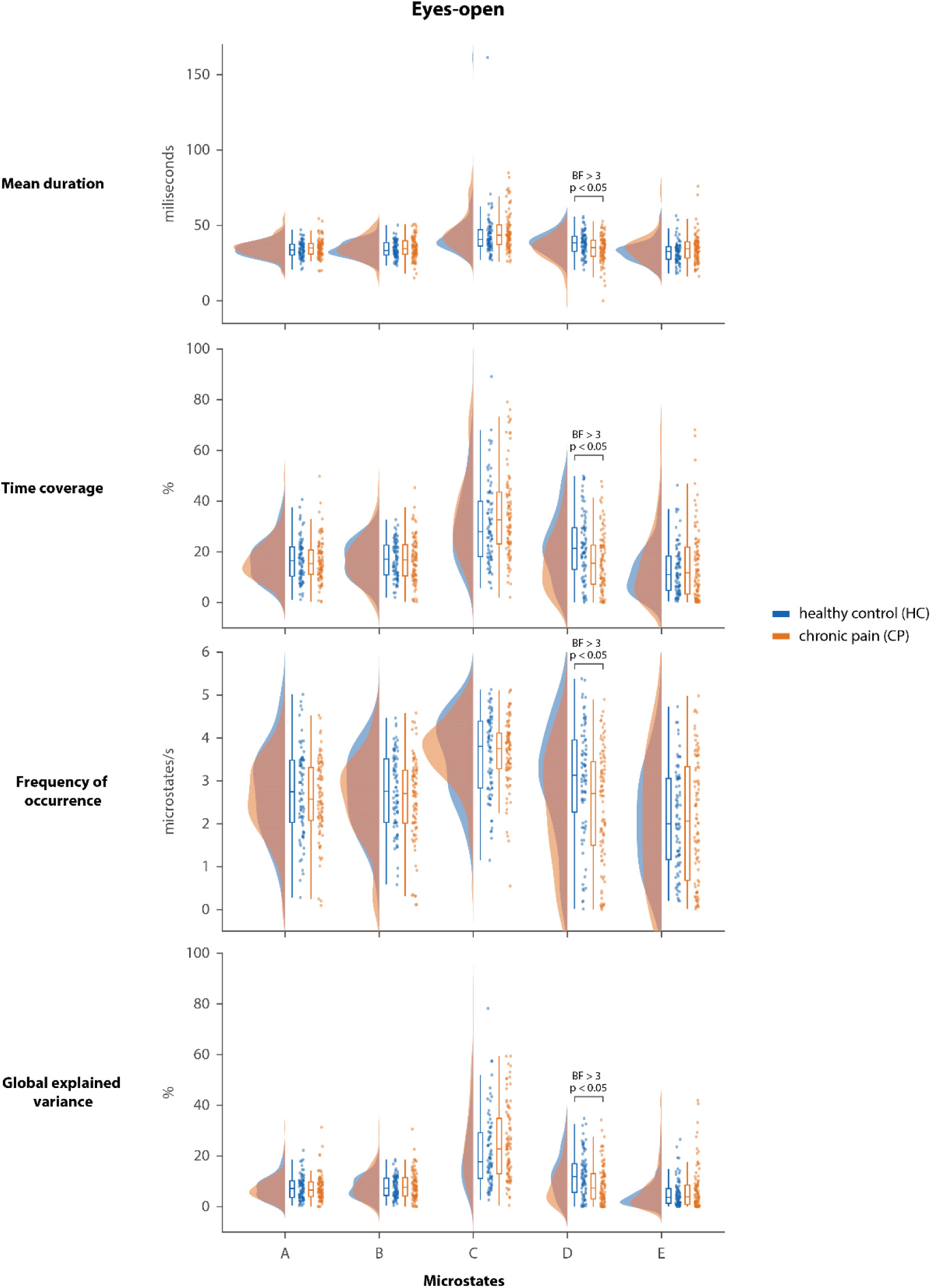
Temporal microstate characteristics in the eyes-open condition. Mean duration, time coverage, frequency of occurrence and global explained variance of each microstate were calculated for chronic pain patients and healthy controls. Raincloud plots [1] show un-mirrored violin plots displaying the probability density function of the data, boxplots, and individual data points. Boxplots depict the sample median as well as first (Q1) and third quartiles (Q3). Whiskers extend from Q1 to the smallest value within Q1 – 1.5* interquartile range (IQR) and from Q3 to the largest values within Q3 + 1.5* IQR. BF = Bayes Factor in favor of the alternative hypothesis.

We further investigated whether the sequences of microstates differed between groups. To this end, transition probabilities from each microstate to all other microstates were calculated for both the eyes-open and the eyes-closed conditions. Mean transition probabilities for both groups and conditions as well as statistical results are presented in tables 4 and 5. We found moderate evidence for a lower transition probability from microstates A and B to microstate D in patients compared to controls in the eyes-open condition (Table 5; 3 < BF_10_ < 10, FDR-corrected p-values = 0.103). Evidence for differences of all other transition probabilities was either inconclusive or provided evidence for no group differences (see tables 4 and 5 for details).

**Table 4.** Group comparisons of transition probabilities between microstates in the eyes-closed condition. Results of two-sided independent samples t-test (frequentist and Bayesian approach). P-values are FDR-corrected. Color coded in red p < 0.05 and BF_10_ > 3, indicating at least moderate evidence for the alternative hypothesis and in blue BF_10_ < 1/3, indicating at least moderate evidence for the null hypothesis. Median effect sizes (δ) and their respective 95% credible interval (CI) are reported. Mean trans. prob. = mean transition probability; CP = chronic pain patients; HC = healthy controls; BF_10_ = Bayes Factor in favor of the alternative hypothesis.

| | Mean trans. prob. CP | Mean trans. prob. HC | t | p | $BF_{10}$ | Median effect size ($\delta$) | 95% CI |
| --- | --- | --- | --- | --- | --- | --- | --- |
| From A to B | 0.232 | 0.221 | -0.595 | 0.920 | 0.188 | -0.081 | [-0.359, 0.196] |
| From A to C | 0.388 | 0.388 | 0.005 | 1.000 | 0.159 | 0.001 | [-0.276, 0.278] |
| From A to D | 0.186 | 0.229 | 2.110 | 0.251 | 1.249 | 0.289 | [0.009, 0.573] |
| From A to E | 0.188 | 0.158 | -1.808 | 0.299 | 0.724 | -0.247 | [-0.530, 0.032] |
| From B to A | 0.223 | 0.224 | 0.073 | 1.000 | 0.160 | 0.010 | [-0.267, 0.287] |
| From B to C | 0.396 | 0.393 | -0.111 | 1.000 | 0.160 | -0.015 | [-0.292, 0.262] |
| From B to D | 0.190 | 0.235 | 2.310 | 0.251 | 1.873 | 0.317 | [0.037, 0.602] |
| From B to E | 0.187 | 0.145 | -2.346 | 0.251 | 2.024 | -0.322 | [-0.607, -0.041] |
| From C to A | 0.232 | 0.251 | 0.923 | 0.811 | 0.237 | 0.126 | [-0.151, 0.405] |
| From C to B | 0.257 | 0.231 | -1.464 | 0.362 | 0.431 | -0.200 | [-0.481, 0.078] |
| From C to D | 0.273 | 0.315 | 1.576 | 0.362 | 0.505 | 0.215 | [-0.063, 0.497] |
| From C to E | 0.235 | 0.201 | -1.739 | 0.299 | 0.648 | -0.238 | [-0.520, 0.041] |
| From D to A | 0.165 | 0.203 | 2.066 | 0.251 | 1.148 | 0.283 | [0.003, 0.567] |
| From D to B | 0.184 | 0.190 | 0.353 | 1.000 | 0.169 | 0.048 | [-0.229, 0.326] |
| From D to C | 0.398 | 0.411 | 0.483 | 0.983 | 0.177 | 0.066 | [-0.211, 0.344] |
| From D to E | 0.201 | 0.182 | -0.848 | 0.828 | 0.222 | -0.115 | [-0.394, 0.161] |
| From E to A | 0.193 | 0.207 | 0.726 | 0.902 | 0.203 | 0.099 | [-0.178, 0.377] |
| From E to B | 0.202 | 0.169 | -1.873 | 0.299 | 0.810 | -0.256 | [-0.539, 0.023] |
| From E to C | 0.384 | 0.368 | -0.648 | 0.920 | 0.193 | -0.088 | [-0.366, 0.189] |
| From E to D | 0.216 | 0.253 | 1.489 | 0.362 | 0.446 | 0.203 | [-0.075, 0.484] |

**Table 5.** Group comparisons of transition probabilities between microstates in the eyes-open condition. Results of two-sided independent samples t-test (frequentist and Bayesian approach). P-values are FDR-corrected. Color coded in red p < 0.05 and BF_10_ > 3, indicating at least moderate evidence for the alternative hypothesis and in blue BF_10_ < 1/3, indicating at least moderate evidence for the null hypothesis. Median effect sizes (δ) and their respective 95% credible interval (CI) are reported. Mean trans. prob. = mean transition probability; CP = chronic pain patients; HC = healthy controls; BF_10_ = Bayes Factor in favor of the alternative hypothesis.

| | Mean trans. prob. CP | Mean trans. prob. HC | t | p | $BF_{10}$ | Median effect size ($\delta$) | 95% CI |
| --- | --- | --- | --- | --- | --- | --- | --- |
| From A to B | 0.236 | 0.245 | 0.598 | 0.810 | 0.196 | 0.085 | [-0.204, 0.376] |
| From A to C | 0.388 | 0.341 | -1.984 | 0.202 | 1.018 | -0.283 | [-0.580, 0.009] |
| From A to D | 0.201 | 0.256 | 2.675 | 0.103 | 4.378 | 0.384 | [0.089, 0.685] |
| From A to E | 0.169 | 0.155 | -0.756 | 0.810 | 0.217 | -0.107 | [-0.399, 0.182] |
| From B to A | 0.244 | 0.243 | -0.097 | 1.000 | 0.167 | -0.014 | [-0.303, 0.276] |
| From B to C | 0.371 | 0.350 | -0.816 | 0.810 | 0.226 | -0.116 | [-0.407, 0.173] |
| From B to D | 0.194 | 0.248 | 2.865 | 0.103 | 7.042 | 0.412 | [0.116, 0.714] |
| From B to E | 0.186 | 0.156 | -1.454 | 0.462 | 0.441 | -0.207 | [-0.501, 0.084] |
| From C to A | 0.272 | 0.249 | -1.234 | 0.609 | 0.336 | -0.175 | [-0.468, 0.115] |
| From C to B | 0.264 | 0.260 | -0.209 | 1.000 | 0.170 | -0.030 | [-0.319, 0.260] |
| From C to D | 0.249 | 0.294 | 2.011 | 0.202 | 1.070 | 0.287 | [-0.005, 0.584] |
| From C to E | 0.209 | 0.194 | -0.719 | 0.810 | 0.211 | -0.102 | [-0.393, 0.187] |
| From D to A | 0.199 | 0.241 | 2.070 | 0.202 | 1.193 | 0.296 | [0.003, 0.593] |
| From D to B | 0.210 | 0.214 | 0.257 | 1.000 | 0.171 | 0.036 | [-0.253, 0.326] |
| From D to C | 0.371 | 0.355 | -0.686 | 0.810 | 0.207 | -0.097 | [-0.388, 0.192] |
| From D to E | 0.204 | 0.186 | -0.804 | 0.810 | 0.224 | -0.114 | [-0.406, 0.175] |
| From E to A | 0.205 | 0.196 | -0.598 | 0.810 | 0.196 | -0.085 | [-0.376, 0.204] |
| From E to B | 0.188 | 0.198 | 0.650 | 0.810 | 0.202 | 0.092 | [-0.197, 0.383] |
| From E to C | 0.386 | 0.342 | -1.921 | 0.202 | 0.909 | -0.274 | [-0.571, 0.018] |
| From E to D | 0.214 | 0.259 | 2.078 | 0.202 | 1.210 | 0.297 | [0.004, 0.594] |

In summary, the analysis of the temporal dynamics of microstates revealed a less predominant role of microstate D in resting-state brain activity of chronic pain patients in both eyes-open and eyes-closed conditions. No evidence for changes of the temporal characteristics of microstates other than D was found.

### 3.3. Relationships between temporal microstate characteristics and clinical characteristics

Having observed consistent evidence for changes of microstate D temporal characteristics in patients with chronic pain, we explored whether microstate D temporal measures were significantly related to clinical characteristics. To this end, we performed correlation analyses between the temporal measures of microstate D and clinical parameters of the patients. We specifically related microstate D characteristics to the current pain intensity, the SF-MPQ total pain score as well as measures of depression (BDI) and medication (MQS). We only found moderate evidence for an inverse relation between medication and the frequency of occurrence of microstate D (Tables 6 and 7; eyes-closed: BF_10_ = 4.16, FDR-corrected p-value = 0.074; eyes-open: BF_10_ = 9.58, FDR-corrected p-value = 0.050). Evidence for an association between medication and the other microstate D measures was inconclusive (Tables 6 and 7; 1/3 < BF_10_ < 3). In addition, evidence for an association between the other clinical characteristics and microstate D measures was either inconclusive or against an association in both conditions (Tables 6 and 7).

**Table 6.** Relationships between microstate D temporal measures and clinical parameters in the eyes-closed condition. Pearson’s correlations (frequentist and Bayesian approach) were performed for microstate D temporal measures, which had consistently shown evidence for differences between patients and controls in previous analyses. P-values are FDR-corrected. Color coded in red p < 0.05 and BF_10_ > 3, indicating at least moderate evidence for the alternative hypothesis and in blue BF_10_ < 1/3, indicating at least moderate evidence for the null hypothesis. ; BF_10_ = Bayes Factor in favor of the alternative hypothesis; 95% CI = 95% credible interval; Mean dur. = mean duration; Time cov. = time coverage; Freq of occ. = frequency of occurrence; GEV = global explained variance; SF-MPQ = short-form McGill pain questionnaire; BDI = Beck’s depression inventory; MQS = medication quantification scale.

|  |  | Current pain | SF-MPQ | BDI | MQS |
| --- | --- | --- | --- | --- | --- |
| Mean dur. | Pearson's r | 0.123 | 0.071 | 0.061 | -0.236 |
|  | p | 0.504 | 0.580 | 0.580 | 0.074 |
| | $BF_{10}$ | 0.260 | 0.160 | 0.150 | 1.974 |
|  | 95% CI | [-0.075, 0.309] | [-0.126, 0.261] | [-0.136, 0.252] | [-0.408, -0.041] |
| Time cov. | Pearson's r | 0.163 | 0.064 | 0.079 | -0.238 |
|  | p | 0.282 | 0.580 | 0.580 | 0.074 |
| | $BF_{10}$ | 0.456 | 0.153 | 0.170 | 2.064 |
|  | 95% CI | [-0.035, 0.345] | [-0.133, 0.255] | [-0.118, 0.269] | [-0.410, -0.043] |
| Freq. of occ. | Pearson's r | 0.236 | 0.074 | 0.056 | -0.265 |
|  | p | 0.074 | 0.580 | 0.580 | 0.074 |
| | $BF_{10}$ | 1.919 | 0.136 | 0.146 | 4.166 |
|  | 95% CI | [0.039, 0.409] | [-0.124, 0.264] | [-0.141, 0.247] | [-0.434, -0.071] |
| GEV | Pearson's r | 0.116 | 0.066 | 0.094 | -0.189 |
|  | p | 0.504 | 0.580 | 0.580 | 0.190 |
| | $BF_{10}$ | 0.240 | 0.155 | 0.193 | 0.718 |
|  | 95% CI | [-0.082, 0.302] | [-0.131, 0.257] | [-0.103, 0.283] | [-0.367, 0.008] |

**Table 7.** Relationships between microstate D temporal measures and clinical parameters in the eyes-open condition. Pearson’s correlations (frequentist and Bayesian approach) were performed for microstate D temporal measures, which had consistently shown evidence for differences between patients and controls in previous analyses. P-values are FDR-corrected. Color coded in red p < 0.05 and BF_10_ > 3, indicating at least moderate evidence for the alternative hypothesis and in blue BF_10_ < 1/3, indicating at least moderate evidence for the null hypothesis. ; BF_10_ = Bayes Factor in favor of the alternative hypothesis; 95% CI = 95% credible interval; Mean dur. = mean duration; Time cov. = time coverage; Freq of occ. = frequency of occurrence; GEV = global explained variance; SF-MPQ = short-form McGill pain questionnaire; BDI = Beck’s depression inventory; MQS = medication quantification scale.

|  |  | Current pain | SF-MPQ | BDI | MQS |
| --- | --- | --- | --- | --- | --- |
| Mean dur. | Pearson's r | 0.191 | 0.108 | 0.177 | -0.164 |
|  | p | 0.279 | 0.434 | 0.279 | 0.279 |
| | $BF_{10}$ | 0.648 | 0.218 | 0.514 | 0.427 |
|  | 95% CI | [-0.018, 0.379] | [-0.101, 0.304] | [-0.032, 0.366] | [-0.354, 0.044] |
| Time cov. | Pearson's r | 0.129 | 0.023 | 0.124 | -0.262 |
|  | p | 0.394 | 0.881 | 0.394 | 0.099 |
| | $BF_{10}$ | 0.272 | 0.136 | 0.256 | 2.861 |
|  | 95% CI | [-0.080, 0.324] | [-0.183, 0.227] | [-0.085, 0.319] | [-0.440, -0.057] |
| Freq. of occ. | Pearson's r | 0.168 | -0.016 | 0.048 | -0.308 |
|  | p | 0.279 | 0.881 | 0.804 | 0.051 |
| | $BF_{10}$ | 0.449 | 0.134 | 0.146 | 9.589 |
|  | 95% CI | [-0.041, 0.358] | [-0.220, 0.190] | [-0.159, 0.250] | [-0.478, -0.105] |
| GEV | Pearson's r | 0.105 | 0.017 | 0.154 | -0.220 |
|  | p | 0.434 | 0.881 | 0.301 | 0.200 |
| | $BF_{10}$ | 0.213 | 0.134 | 0.366 | 1.106 |
|  | 95% CI | [-0.104, 0.302] | [-0.189, 0.221] | [-0.056, 0.345] | [-0.402, -0.012] |

Taken together, the results did not provide consistent evidence for relationships between microstate D temporal measures and clinical characteristics.

## 4. Discussion

In the present study, we investigated the dynamics of brain function in patients suffering from chronic pain. To this end, we performed microstate analysis of resting-state EEG recordings in a large cohort of patients and age- and sex-matched healthy control participants. In both groups, resting-state brain activity could be described as sequences of 5 microstates labeled A to E. Analyses of the temporal characteristics of these microstates revealed a decreased presence of microstate D in patients as compared to healthy participants. These changes were consistently found in eyes-open and eyes-closed conditions. No evidence for differences in other microstates was found. Thus, the present findings describe microstate D-specific changes of the dynamics of brain function in resting-state EEG recordings of patients suffering from chronic pain.

Our analyses reveal shorter mean duration, lower time coverage, fewer occurrences and less explained variance of microstate D in patients compared to controls in both eyes-open and eyes-closed recordings. Thus, our findings robustly indicate a decreased presence of microstate D in resting-state EEG activity in patients suffering from chronic pain. These findings are in contrast to the only study which applied microstate analysis to resting-state EEG recordings of patients suffering from chronic pain so far [20]. That study found a lower occurrence and time coverage of microstate C in eyes-open resting state recordings of 43 patients. This difference between studies might at least in part be due to differences between patient groups. The previous study included 43 patients suffering from fibromyalgia whereas the present study included 101 patients suffering from different chronic pain conditions. Although the heterogeneous sample of the present study reflects clinical reality it might veil differences between chronic pain conditions. The two studies together might prompt further studies in larger groups of patients, ideally from different recording sites, to resolve this issue and to further clarify changes of microstates in chronic pain.

Our observations further complement recent fMRI studies, which have shown changes of the dynamics of brain function in chronic pain at ultra-low frequencies below 0.1 Hz (e.g. [3; 5; 10]). They extend this evidence by showing alterations of the dynamics of brain function at frequencies higher than 1 Hz, in line with the *dynamic pain connectome* concept [28; 45].

Microstate analysis is an emerging tool for investigating the dynamics of brain activity. Although the functional interpretation of microstates is not fully clear yet, microstate analysis has been increasingly used to identify changes of brain dynamics in various neuropsychiatric diseases [25; 34], which have furthered the understanding of the pathology of these disorders. Beyond, alterations of microstates’ characteristics might be useful as clinical biomarkers. For instance, a recent study has identified the dynamics of microstates C and D as a promising candidate endophenotype for schizophrenia [12]. However, our data did not provide evidence for a correlation between alterations of microstate D and clinical characteristics. As the brain processes discriminating chronic pain patients from healthy people differ from those encoding momentary pain intensity [3; 32; 54; 58; 63], the observed changes might reflect the abnormal disease state *per se* rather than its specific characteristics.

Microstate D has been related to attentional brain networks and functions (for reviews see [34; 51]). In particular, microstate D has been associated with brain activity in fronto-parietal regions [8; 11], the dorsal attentional control network [49] and focus-switching and attentional reorientation [36]. Interestingly, deficits of cognitive function and particularly of attentional switching have been extensively reported in patients suffering from chronic pain [37]. A common hypothesis is that pain competes with other stimuli for limited cognitive resources, thereby “demanding attention” and potentially impairing higher-order attentional control mechanisms [17; 30; 56]. Thus, a decreased presence of microstate D might represent a neurophysiological correlate of altered attentional functioning in chronic pain. However, as we have not obtained direct measures of attentional functioning, we cannot directly test this hypothesis in the present study. Future microstate studies on chronic pain might therefore include tasks and/or questionnaires assessing attentional functions.

Several limitations of the current study need to be discussed. First, the specificity of the decreased presence of microstate D for chronic pain is unclear. In particular, studies in patients suffering from schizophrenia [12; 47] and major depressive disorder [38] also showed a decreased presence of microstate D. However, investigating symptom- and disease-specificity of these findings is challenging. Substantial progress in this endeavor requires large samples of patients suffering from different neuropsychiatric symptoms and diseases, standardized assessments and, ideally, sharing of data acquired at different sites. As a first step in that direction, we share data and code of the present study in a standardized format with the research community. Second, comparisons of microstate topographies showed slight but statistically significant differences between patients and healthy controls. However, considering these subtle differences together with the overwhelming similarity of microstate topographies, the microstates of both groups likely capture the same underlying neural networks. Third, the causal relationship between altered microstate dynamics and chronic pain is unclear. First studies have shown that the dynamics of microstates can be changed by neurofeedback [15] and non-invasive brain stimulation [53]. These approaches might thus be useful to prove the causal link between changes in microstate dynamics and neuropsychiatric disorders including chronic pain. Moreover, they highlight the potential utility of microstate dynamics as targets for neurofeedback- and/or brain stimulation-based treatments of chronic pain.

In conclusion, our findings provide evidence for altered dynamics of brain function in a large cohort of chronic pain patients using EEG microstate analysis. We particularly observed alterations of microstate D. As this microstate has been associated with attentional brain networks and functions, changes of microstate D might relate to dysfunctional attentional processes in chronic pain. These results add to the understanding of the pathophysiology of chronic pain. Moreover, they might be useful for establishing a much sought-after EEG-based biomarker of chronic pain.

## Acknowledgments

The study was supported by the Deutsche Forschungsgemeinschaft (PL 321/10-2, PL321/11-2, PL321/13-1). The authors have no conflict of interest to declare.

